# Cortical spheroids display oscillatory network dynamics

**DOI:** 10.1101/2021.07.18.452530

**Authors:** Jessica L. Sevetson, Brian Theyel, Diane Hoffman-Kim

**Author notes:** Corresponding Author, Electronic Supplementary Information (ESI) available.

## Abstract

Three-dimensional brain cultures can facilitate the study of central nervous system function and disease, and one of the most important components that they present is neuronal activity on a network level. Here we demonstrate network activity in rodent cortical spheroids while maintaining the networks intact in their 3D state. Networks developed by nine days in culture and became more complex over time. To measure network activity, we imaged neurons in rat and mouse spheroids labelled with a calcium indicator dye, and in mouse spheroids expressing GCaMP. Network activity was evident when we electrically stimulated spheroids, was abolished with glutamatergic blockade, and was altered by GABAergic blockade or partial glutamatergic blockade. We quantified correlations and distances between somas with micron-scale spatial resolution. Spheroids seeded at as few as 4,000 cells gave rise to emergent network events, including oscillations. These results are the first demonstration that self-assembled rat and mouse spheroids exhibit network activity consistent with in vivo network events. These results open the door to experiments on neuronal networks that require fewer animals and enable high throughput experiments on network-perturbing alterations in neurons and glia.

## Introduction

There is a widely acknowledged need for models of the brain that present *in vivo* cell types as well as network activity, and allow high throughput assessment of the effects that various perturbations have on such network activity. Developments in three-dimensional culturing methods over the past decade ^1–5^ have facilitated the creation of *in vitro* platforms with increasing *in vivo* relevance. While much research has been done to observe and characterize these three-dimensional platforms, for example, at the transcriptomic level (for review, see Amin & Paşca, 2018; Gopalakrishnan, 2019; Kratochvil et al., 2019; Marton & Paşca, 2020), multiple factors limit their use. These include a lack of reproducibility of size and composition (though some progress has been made; see Velasco et al., 2019), the technical complexity of integrating non-neural cell types ^11,12^, and the long timescale required for development of network activity ^13^.

Understanding neuronal network activity *in vitro* is fundamental to our ability to use three-dimensional neural tissues to address key questions about development and disease. Electrical activity in the brain is ubiquitous in vertebrates and is necessary for healthy human brain function. Synchronous neuronal firing is a key feature of developing neural networks *in vivo* ^14–19^. Neural network activity evolves in a robust, stereotyped fashion: sparse, uncorrelated activity is followed by highly synchronized oscillatory activity, which then develops into the complex network activity consistent with mature phenotypes. Many injuries, diseases, and neurodevelopmental disorders display deficits at the network level, some of which have been observed in culture. For example, cells derived from the iPSCs of autistic patients exhibited decreased network connectivity compared to cultures derived from the iPSCs of non-autistic subjects ^7,8,20–22^.

In previous work, we have shown that spheroids derived from rodent cortex contain multiple cell populations – including astrocytes, microglia, endothelial cells, and active neurons – at physiological cell density and tissue stiffness ^3,23^. Synaptic activity was confirmed with patch clamp of individual neurons. Still unknown, however, was to what extent the neurons within a spheroid form spontaneous, active networks.

Here, we report the results of our cross-sectional observations of the network activity in cortical spheroids from rat and mouse. We employed calcium imaging to noninvasively observe network-level neuronal activity. We characterized activity across the development of the spheroid and found that synaptically driven network activity developed in a reproducible manner and was dependent on seeding density. We showed quantitatively that activity was similar in cortical spheroids derived from male and female rats We also showed qualitatively that activity was similar between rats and mice. We have identified network-level activity in cortical spheroids with similarities to activity observed *in vivo*. This suggests that cortical spheroids could be used in a high-throughput way to determine what potential effects disease modeling and therapeutic strategies have on network activity.

## Experimental Procedures

### Animal usage

All animal procedures were conducted in accordance with NIH guidelines. Procedures were approved by Brown University’s Institutional Animal Care and Use Committee. For studies using rats, CD rats (Charles River) were used. For studies using endogenously fluorescent mice, cre expression was driven via the Retinol Binding Protein promoter, crossed with lox-GCaMP6f (Jackson Labs, #003967) to enable expression of fluorescent calcium indicator in RBP-positive pyramidal neurons (B6.Cg-Tg(Rbp3-cre)528Jxm/J; gift of Dr. Chris Moore). Sex in perinatal rat and mouse pups was determined by angiogenital distance.

### Rat spheroid preparation

Spheroid preparation proceeded as described in a previous study (Dingle et al., 2015). In brief, cortex was isolated from male or female rats (Charles River) at postnatal Day 0-2, diced, and enzymatically digested in 2 mg/mL solution of papain (BrainBits, LLC) in Hibernate A sans Ca^2+^ (BrainBits, LLC) for 30 minutes at 30°C. The papain solution was removed, and cells were mechanically separated with a fire-polished Pasteur pipette in Hibernate A buffer supplemented with 1 × B27 (Invitrogen) and 0.5 mM GlutaMAX (Invitrogen). The cell solution was then centrifuged and resuspended in cortical media consisting of Neurobasal A (Invitrogen) supplemented with B27, 0.5 mM GlutaMAX, and Penicillin-Streptomycin (Invitrogen). The cell suspension was washed again by centrifuging and resuspending the pellet in cortical media. Cortical cells were placed in 2% agarose microarrays created by 400 μm diameter round silicon peg molds (Microtissues, Inc) at a density of 4-8,000 cells/microwell. The cell suspension was allowed to settle in the microwells for 30-40 minutes, after which 1 mL of cortical media was added to each well of a 24-well plate. Spheroids were incubated at 37° C in 5% CO_2_, and the cortical media was changed every 2-3 days.

### Mouse GCaMP6f spheroids

Mouse cortex was dissected and dissociated at P0-3. Dissection proceeded identically to rat pup dissection, except that the tissues were only incubated in papain for 15 minutes. As mouse pups were too young to genotype prior to dissection, pairs of cortices were pooled together to ensure the greatest number of samples with fluorescent neurons. Anogenital distance was used to determine sex, and cortices of the same sex were pooled together.

### Calcium dye imaging

Rat spheroids were incubated with 5.2 μM Oregon Green 488 BAPTA-1 AM (OGB-1, ThermoFisher) and 0.02% Pluronic acid (ThermoFisher) in cortical media, in the dark for 20-25 minutes at 37°C and 5% CO_2_. After incubation, the well was washed twice with media to remove excess dye. Spheroids were removed from the agarose gel by gently pipetting up and down over the wells, then transferred to a 35 mm petri dish with a glass bottom for confocal imaging. Images were acquired using a high sensitivity resonant scanner in an Olympus FV3000-RS with HV from 400-500, gain of 1, and offset of 6. Spheroids labeled with OGB were imaged with 0.6% laser intensity, and spheroids labeled with GCaMP6f were imaged with 2% laser intensity to compensate for the comparative dimness of genetically encoded calcium indicators. Cells were kept at 37° C throughout the imaging session and were imaged at 25-50 FPS for 1 minute unless otherwise noted. Images were converted to .tif format, and Z-stacks of 50-150 μm were collapsed into a single plane for viewing using Fiji ImageJ software.

### Pharmacology

NBQX (Tocris 1044) was used at a concentration of 20 μM or 2 μM and incubated with the spheroid for 10 minutes prior to imaging. AP-5 (Tocris 0105) was used at a concentration of 50 μM or 5 μM and incubated with the spheroid for 10 minutes prior to imaging. Bicuculline (BCC; Sigma 14340) was used at 10 μM and incubated with the spheroid for 20 minutes prior to imaging.

### Electrophysiology

Female-derived spheroids of approximately 4,000 (4K) or 8,000 (8K) cells each were allowed to adhere for 48 hours to glass coverslips coated in 50 mM/mL Poly-d-lysine (PDL; Sigma). Spheroids were transferred to a submersion recording chamber, where they were bathed continuously (3-4 ml/min) with warm (32°C) oxygenated (95% O2,5% CO2) ACSF containing 126 mM NaCl, 3 mM KCl, 1.25 mM NaH2PO4, 1 mM MgSO4, 1.2 mM CaCl2, 26 mM NaHCO3, and 10 mM glucose. Cells were visualized with infrared differential interference contrast microscopy. Glass recording pipettes were pulled to final tip resistances between 4 and 7 MW. For current-clamp recordings, micropipettes were filled with internal solution of the following composition (in mM): 130 K gluconate, 4 KCl, 2 NaCl, 10 HEPES, 0.2 EGTA, 4 ATP-Mg, 0.3 GTP- Na, and 14 phosphocreatine-2K. Electrophysiological signals, acquired in current clamp, were amplified with a Multiclamp 700B (Axon), in which signals were first filtered (DC–10 kHz) and then digitized at 20 kHz with the Digidata 1440A data acquisition system and Clampex data acquisition software (Axon).

### Image analysis

Image stacks were converted to TIFF format in ImageJ, and the rest of the analysis proceeded in custom MatLAB software (version 2019b, https://github.com/Jess7son/Ca_imaging_public). Briefly, image stacks were registered, and regions of interest (ROIs) were manually designated based on visible cell borders. Additional ROIs were circled to monitor whole-sphere activity and to check for background noise. Fluorescence over time was extracted and normalized to the first 500 frames for each ROI, and saved with corresponding ROI topographical information as well as metadata regarding spheroid Day in culture, size, sex, pharmacological reagents, etc. Power spectra and primary frequency were calculated from raw ΔF/F traces using the mscohere function (MATLAB, Chronux Toolbox v2.12). Binary active/inactive designations were made based on which cells displayed increased activity (at least a 10% intensification) between 0.05 Hz and 1 Hz as determined by the individual cellular power spectra (See Statistical Analyses). Fluorescence traces were de-noised with a 0.5 second gaussian filter prior to correlation calculations, amplitude calculations, and individual trace plotting. For individual traces, representative active ROIs were manually selected, unless the sample contained <4 active ROIs, in which case some inactive ROIs were plotted.

### Statistical analyses

To detect potential differences in variance between groups, Levene’s Tests were conducted using vartestn (MATLAB Signal Processing Toolbox) with test type set as ‘LeveneAbsolute’ and sensitivity α=0.05. Welch’s ANOVAS with Games-Howell *post hoc* tests were calculated using the wanova and GHtests functions respectively (MATLAB file exchange, Andrew Penn and Antonio Trujillo-Ortiz respectively, both version 1.0.0.0) with sensitivity α=0.01. T-tests (ttest2; MATLAB Statistics and Machine Learning Toolbox) were calculated with sensitivity α=0.01 unless otherwise noted.

## Results & discussion

### Development of network activity across time

Previous studies have demonstrated action potentials and active synapses in 8,000-cell (8K) spheroids by Day 14 ^3^. To investigate the presence of network activity, we created spheroids from rat cortex using previously established self-assembly methods ^3,23^ and incubated them with the calcium indicator Oregon Green 488 BAPTA 1-AM (OGB-1; Fisher). Based on the known timeline in 8K spheroids, wherein action potentials are largely absent at Day 7 but are present both evoked and spontaneously by Day 14, we selected Days 3, 7, 9, and 14 for calcium imaging analysis. In each dissection, cortical tissue was harvested from 2 P0-2 rat pups from the same litter and of the same sex. At each timepoint, six to eight 8K spheroids each from male and female rat cortex were imaged. There was no detectable activity on Day 3 (data not shown). At Days 7, 9, and 14, synchronized activity was readily apparent in the <1Hz range (Fig 1A). Spheroids comprised of 2,000 cells (2K spheroids) and 4,000 cells (4K spheroids) were subjected to the same cross-sectional analysis (Fig S1A-B). Although 2K spheroids never exhibited detectable activity (Fig S1A), some 4K spheroids showed oscillatory activity by Day 21 (Fig S1B-C).

**Figure 1).**
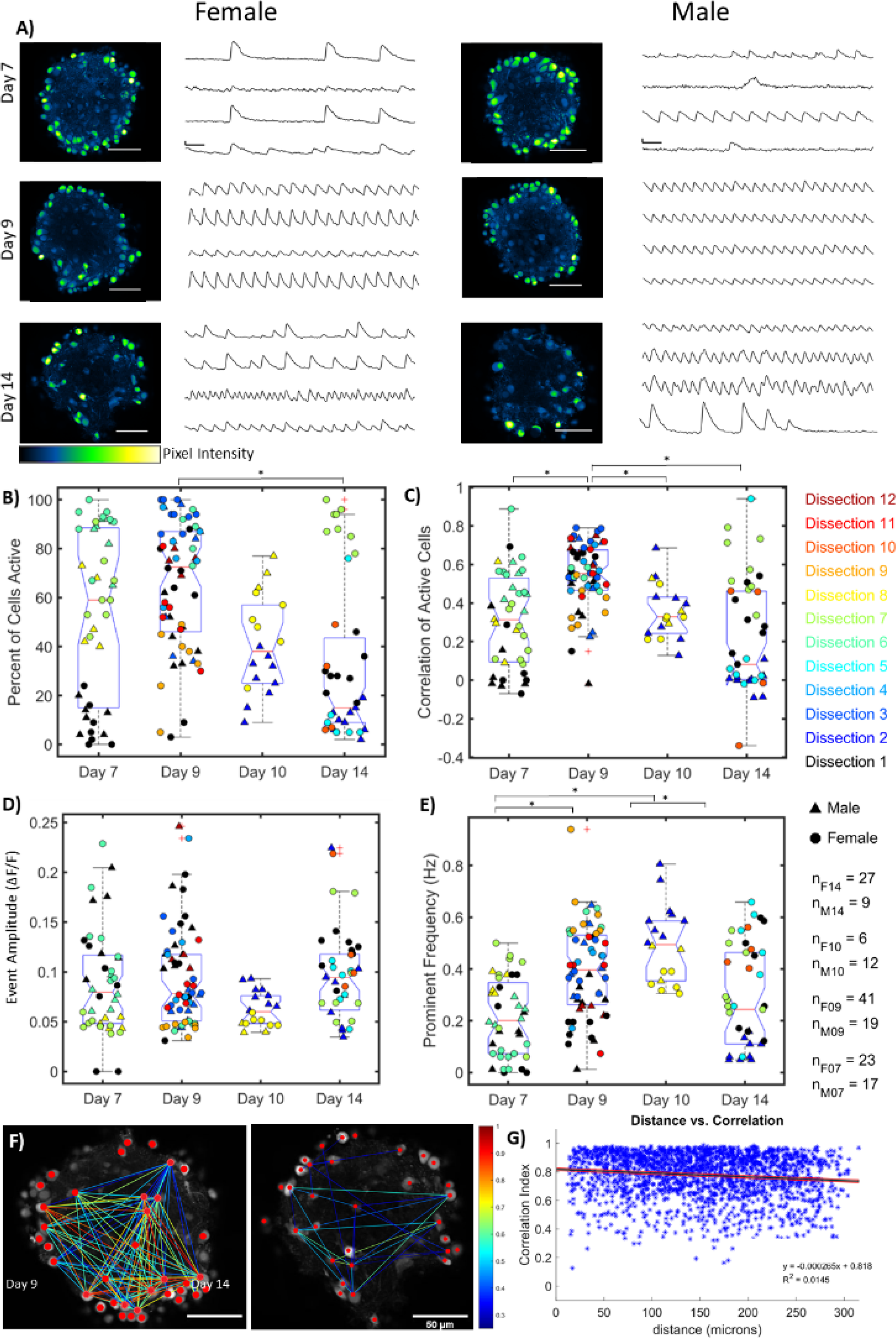
Postnatal rat cortical spheroids displayed network activity that develops over time. A) Confocal images of spheroids labeled with OGB-1 paired with ΔF/F traces from representative neurons. 8K spheroids displayed variable or nonexistent activity at Day 7 (top), oscillatory activity at Day 9 (center) and complex activity at Day 14 (bottom). Image scale bars 50 μM. Time series scale bars are 5 sec (horizontal) by 0.1 ΔF/F (vertical). Activity was not dependent on whether spheroids were derived from female (left column) or male (right column) rat cortex. B-E) Percent of cells active (B), correlation of active cells (C), amplitude of events (D), and frequency of events (E) were compared for spheroids imaged at Day 7 (n = 40), Day 9 (n = 27), Day 10 (n = 38) and Day 14 (n = 35). Marker symbol indicates the sex of rodent the spheroid was derived from (▲=♂, ●=♀). Spheroids from the initial dissection in black, with subsequent dissections denoted in chronological order via marker colors ranging from violet to maroon. At right, n subscripts indicate sex and time in vitro, such that nF07 indicates the total number of Day 07 female spheroids observed. Red + symbols indicate statistically detected outliers. Sample variance was consistent between male- and female-derived tissues in all categories (Levene’s test, p>0.05), pooled across all days in culture. Therefore, unpaired t-tests were conducted and found no significant differences between means of male- and female-derived spheroids in any category with α=0.01. As sample size and variance differed in all categories between timepoints (Levene’s test, p<0.05), Welch’s ANOVA was used with Games-Howell pairwise comparisons to compare group means with α=0.01. B) Spheroids contained more active cells at Day 9 than at Day 14 (μ7 = 53% ± 34%, μ9 = 66% ± 26%, μ10 = 40% ± 19%, μ14 = 35% ± 33%). C) Calcium activity in spheroids was most highly correlated at Day 9 (μ7 = 0.31 ± 0.24, μ9 = 0.54 ± 0.17, μ10 = 0.36 ± 0.14, μ14 = 0.25 ± 0.29). D) Amplitude of calcium events in spheroids was consistent over time (μ7 = 0.089 ± 0.052, μ9 = 0.093 ± 0.048, μ10 = 0.063 ± 0.017, μ14 = 0.10 ± 0.045). E) The prevalent frequency was lower at Day 7 than Days 9 or 10. The prevalent frequency was higher at Day 10 than at Day 14 (μ7 = 0.21 ± 0.15 Hz, μ9 = 0.39 ± 0.17 Hz, μ10 = 0.50 ± 0.14 Hz, μ14 = 0.31 ± 0.19 Hz). F) Spheroids were more correlated at Day 9 than at Day 14. Sample correlation diagrams overlaying confocal images. Diagrams show cell somas (red dots) connected by colored lines indicating correlation strength between the two cells. Correlation strengths range from .25 (blue) to 1 (maroon). Correlation was higher at Day 9 (left) than at Day 14 (right), and correlated cells were distributed throughout the plane of view. G) Distance within the plane of view had no noticeable effect on correlation strength. Sample linear regression model (black line) and 99% confidence interval (red lines). Correlation vs in-plane distance is plotted between all pairs of cells (blue stars) from a Day 9 Spheroid. Slope = −.0265% correlation per micron, R2 = 0.0145.

For timepoints at which activity was observed, the imaging protocol was replicated to account for potential batch effects, and expanded to include Day 10. Cells within each spheroid exhibiting a >10% increase in average power in the 0.1 – 1 Hz frequency band were declared ‘active’ and selected for further analysis. Spheroids were compared based on the proportion of active cells (Fig 1B), the average correlation between those cells (Fig 1C), the average event amplitude (Fig 1D), and the most prevalent frequency of activity (Fig 1E). Spheroids derived from female and male cortex exhibited similar patterns of activity (Fig 1A-E; n_Male_=57, n_Female_=97), with no significant difference between variances and means (Levene’s test p>0.05, unpaired t-test p>0.01). As the variances between timepoints were unequal (Levene’s test, p>0.05) and sample sizes were also unequal, we used Welch’s ANOVA with Games-Howell *post hoc* tests (α=0.01) as a more conservative determinant of significance.

At Day 7 (4 imaging replicates, n=40), the proportion of cells within a plane of view exhibiting dynamic activity was variable (Fig 1B). Activity, when present, was generally low in both correlation (Fig 1C) and in frequency (Fig 1E). By Day 9, spheroids consistently showed oscillatory activity (6 replicates, n=60). This activity tended to involve most cells in view (Fig 1B), was highly synchronized (Fig 1C), and was of a slightly higher frequency (Fig 1E). At Day 10 (2 replicates, n=18), activity began to become less correlated (Fig 1C), with fewer cells being active (Fig 1B). Activity was at a slightly higher frequency than at Day 7 or 14 (Fig 1E). By Day 14 (5 replicates, n=38), activity was again variable in both proportion of cells active (Fig 1B) and in the correlation of those cells (Fig 1C), with some cells firing in phase but at irregular intervals (Fig 1A, Day 14). Event amplitude did not differ across timepoints (Fig 1D).

We asked whether distance between cells within a spheroid had any effect on correlation, and observed that spheroids were more correlated at Day 9 than at Day 14 (Fig 1F). When correlation was plotted as a function of distance in the plane of view (Fig 1G, Day 9), there was no evidence of a relationship between the two. Across all spheres in all conditions, in-plane distance between a given pair of cells only accounted for 2.06% ± 2.59% of variance in correlation (fitted linear regression models of distance vs. correlation in 201 spheres).

### Functional underpinnings of calcium activity

To confirm that the observed calcium fluctuations were the result of electrical and synaptic activity, we used an extracellular electrode to deliver 10 Hz stimulation during calcium imaging. Increases in fluorescence were coincident with electrical stimulation (Fig 2A).

**Figure 2).**
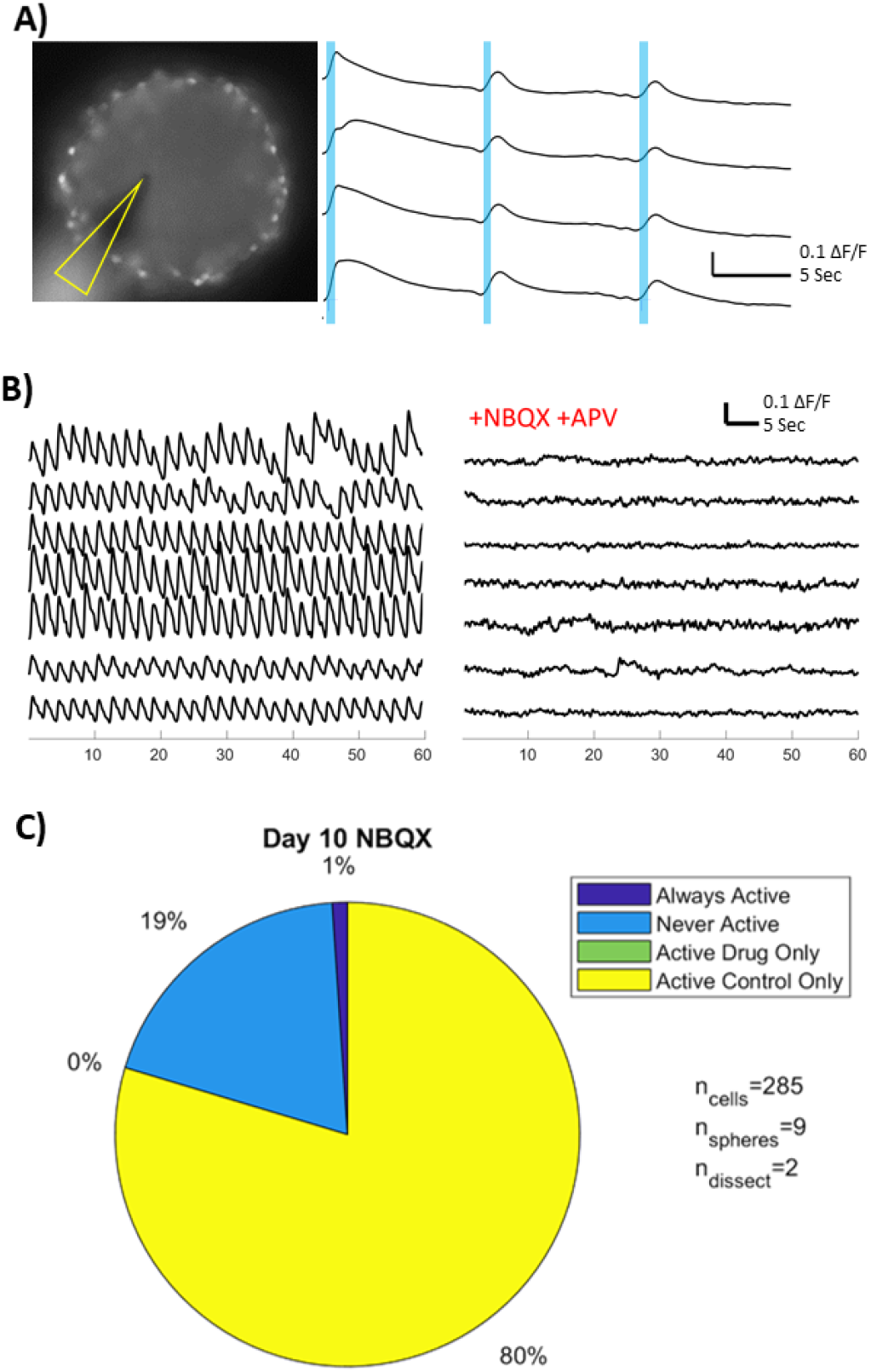
Electrical stimulation was sufficient and excitatory synaptic input was necessary for the presence of network activity in rat spheroids. A) Epifluorescent image of a 4K spheroid at Day 21 labeled with OGB-1 paired with ΔF/F traces of representative neurons. Electrode outlined in yellow. Blue lines indicate stimulation. B-C) Glutamatergic blockade with NBQX and APV abolished dynamic calcium activity. B) ΔF/F traces of representative neurons from a Day 10 spheroid. Break indicates 10-minute bath application. C) Active cells ceased to display dynamic calcium activity following glutamatergic blockade with NBQX/APV. Pie chart depicting proportions of cells that displayed calcium activity only during the control condition (yellow, 80%) or never displayed dynamic calcium activity (blue, 19%). In both dissections, a single cell out of all spheres examined exhibited some activity in both control and glutamatergic blockade conditions (violet, <1%). No cells displayed activity solely under glutamatergic blockade. Results were taken from 285 cells across 9 spheres from 2 separate dissections.

To evaluate whether dynamic fluorescence changes were driven by synaptic activity, we exposed spheroids to 20 μM NBQX and 50 μM AP-V, antagonists of AMPA- and NMDA-type glutamate receptors, respectively. Almost all cells ceased to exhibit calcium activity following glutamatergic blockade (Fig 2B-C), demonstrating that calcium fluorescence changes are dependent upon the presence of excitatory post-synaptic currents.

### Activity in GCaMP6f expressing mouse tissues

The ability to create spheroids out of mouse cortex would allow the model to leverage the powerful genetic tools available in mice. Thus, we investigated whether these patterns were also present in mice. For this experiment, rat Day 9 was selected as the comparison timepoint, as its highly synchronized oscillatory phenotype was particularly robust.

Spheroids created from RBP-cre x GCaMP6f mice contained many cells that exhibited spontaneous fluorescence changes, suggesting successful expression. To account for differences in developmental timing 24,25, mouse spheroids were imaged at Day 8 and Day 13 (Fig 3A). A small number of transgenic mouse spheroids were also incubated with OGB-1 (Fig 3B) to evaluate any indicator-specific differences. The changes in fluorescence were of higher amplitude using GCaMP6f than OGB-1, likely due to the low baseline fluorescence of GCaMP6 indicators ^26^. Mouse-derived spheroids also responded to electrical stimulation, with repeated stimulation amplitude leading to additional neuronal recruitment (Fig 3C). In one RBP-Cre x GCaMP6f mouse-derived spheroid, a 600 ms square-wave pulse of 100 pA triggered some local recruitment, then a second pulse of 75 pA, delivered 10 seconds after the first, led to greater activity and more neurons being recruited (Fig 3C). RBP^+^ neurons within mouse-derived spheroids tended to be active and exhibited high intercellular correlation and low firing frequency (Fig 3D-F).

**Figure 3).**
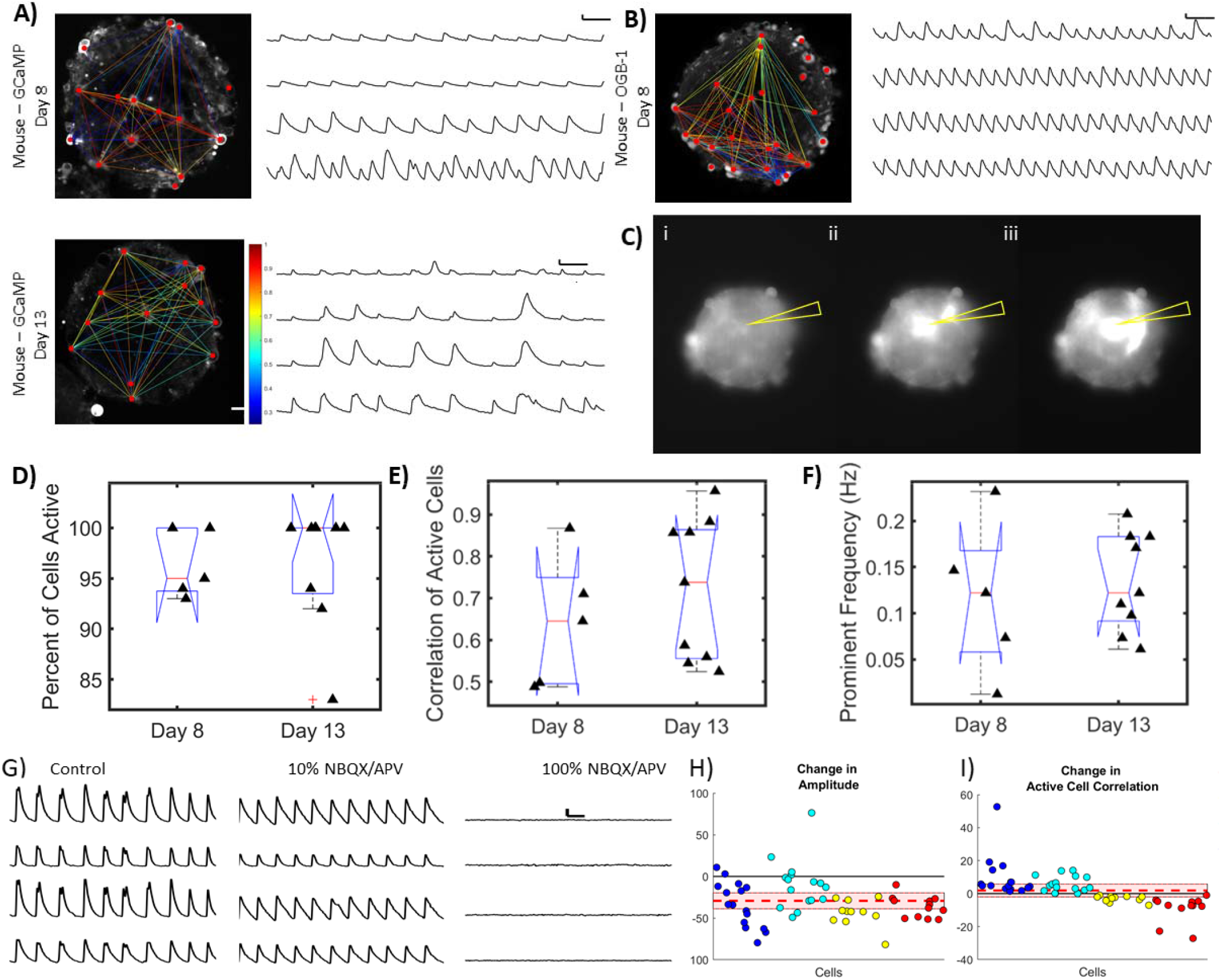
Spheroids developed oscillations within the first two weeks of culture in mouse-derived spheroids that were similar to oscillations previously noted in rat-derived spheroids. Oscillations were observed with genetic as well as chemical calcium indicators. A) Left: Sample correlation diagram overlaying confocal images of mouse derived spheroids (8K spheroid) at Day 8 and 13. Spheroids expressed the genetically encoded GCaMP6f indicator in RBP-expressing pyramidal neurons. Diagram shows cell centers (red dots) connected by colored lines indicating correl tion strength between those two cells. Correlation strengths ranged from .25 (blue) to 1 (maroon). Right: paired ΔF/F traces of representative neurons. Trace scale bars 5 sec (horizontal) by 0.5 ΔF/F B) Spheroids labeled with OGB-1 displayed similar activity patterns as seen with GCaMP6f. Same plots as in A, but in a mouse-derived, 8K spheroid at Day 8 labelled with OGB-1. Trace scale bars 5 sec (horizontal) by 0.1 ΔF/F (vertical; axis adjusted from GCaMP6f for visualization). C) Electrical stimulation was sufficient to activate GCaMP6f in spheroids in an intensity-dependent manner. Epifluorescent images of a mouse-derived, 4K spheroid at Day 21. Spheroid imaged at baseline (i), following the first, 100 pA stimulation (ii) and following the second, 75 pA stimulation (iii) of a single cell in the juxtacellular configuration. Yellow arrow indicates the location of the stimulating electrode. D-F) Percent of cells with detected activity (D), correlation values (E), and prominent frequencies (F) for labeled mouse spheroids at Days 8 and 13. Neither variance (Levene’s Test, p>0.05) nor mean values (unpaired t-test, p>0.01) differed between the groups. G) Activity continued during the 2μM NBQX/5μM APV addition (10%), and was abolished in the presence of 20μM NBQX/50μM APV (100%). Paired ΔF/F traces of cells at baseline (left), with 2 μM NBQX + 5 μM AP-V (10%; center), and with 20 μM NBQX + 50 μM AP-V (100%; right). H) Amplitude decreased following 2μM NBQX/5μM APV addition (t-test, p<0.01). Cell-by-cell data showing percent change in mean event amplitude following 2μM NBQX/5μM APV addition. Event amplitudes were calculated for active cells. Marker color indicates cells from the same spheroid. Red dashed line indicates new mean amplitude, with 99% confidence interval shaded in on either side. Cells experienced a 29% ± 26% decrease in mean event amplitude. I) Intercellular correlation increased in some cells at 2μM NBQX/5μM APV, though this trend was not significant (p>0.01). Cell-by-cell data showing percent change in mean intercellular correlation following 2μM NBQX/5μM APV addition. Intercellular correlations were calculated between active cells. Marker color indicates cells from the same spheroid. Red dashed line indicates new mean correlation, with 99% confidence interval shaded in on either side.

To determine whether excitatory neurotransmission played a role also in mouse tissue, we investigated the effects of decreasing glutamatergic input. The addition of 10% NBQX-APV (2 mM/5 mM respectively) led to decreases in mean event amplitude but no significant change in correlation (Fig 3G-I). Activity in these same spheroids was completely abolished when glutamatergic receptor antagonists were increased to full concentrations (20 mM NBQX/50 mM APV; Fig 3G). The clear cessation of activity after glutamatergic receptor blockade confirmed the presence of glutamatergic synapses in the network, and validated that oscillatory activity was dependent on activity at those synapses.

### Inhibitory contributions to network properties

We next investigated the network contribution of inhibitory synapses. By recording from the same cells before and after GABAergic blockade with bicuculline, we saw that rat spheroids at Day 10 showed a modest phenotype, (Fig 4A). Individual cells rarely switched from active to inactive or vice-versa (Fig 4B) but did display a decreased mean event amplitude and a slightly increased mean event decay time (Fig 4C). Event rise time and intercellular correlation were unaffected. In contrast, GABAergic blockade in spheroids at Day 14 led to a highly synchronized, slow oscillatory pattern of behavior similar to baseline Day 9 activity (Fig 4D). A greater proportion of inactive cells became active, and more active cells became inactive (Fig 4E). While mean event rise time decreased, mean event decay time increased, and event amplitude itself was unchanged (Fig 4F). Inter-cellular correlation dramatically increased (Fig 4F).

**Figure 4).**
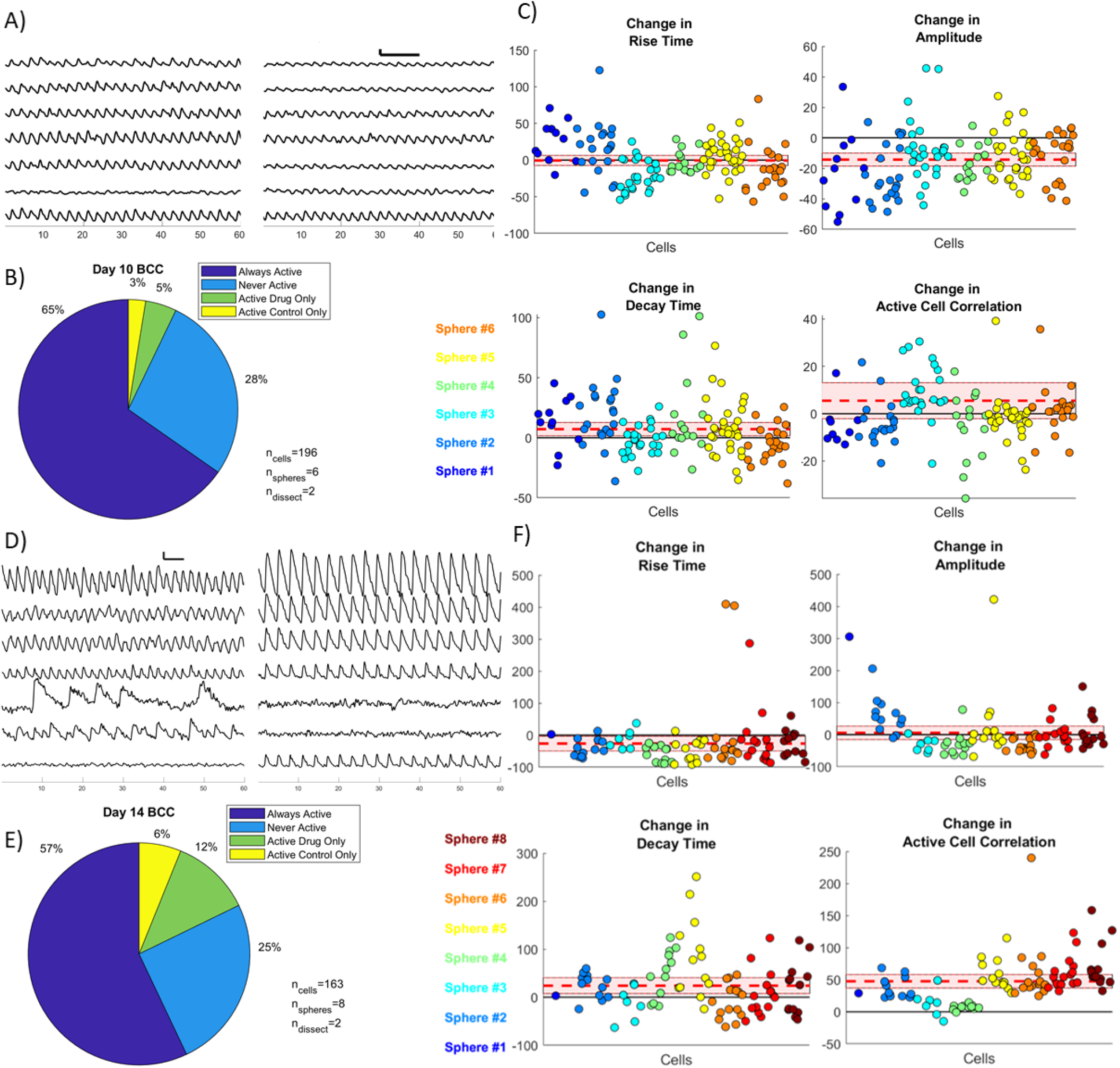
Inhibitory synapses contributed to spheroid networks differently at different stages. A) Amplitude decreased in some cells following GABAergic block with bicuculline in a Day 10 spheroid. ΔF/F traces of cells in an 8K rat spheroid at baseline (left) and after GABAergic block with bicuculline (right). Break represents 20 minute bicuculline incubation. Time series scale bars are 5 sec (horizontal) by 0.1 ΔF/F (vertical). B) Cells rarely switched their activity status following GABAergic blockade with bicuculline at Day 10. Pie chart depicting proportions of cells that exhibited activity in both control and GABAergic blockade (violet, 65%) or never displayed dynamic calcium activity (blue, 28%). Small numbers of cells displayed activity solely under GABAergic blockade (green, 5%) or in the control condition (yellow, 3%). Results were taken from 196 cells across 6 spheres from 2 separate dissections. C) At Day 10, GABAergic blockade via bicuculline incubation yielded a decrease in amplitude and an increase in decay time. Normalized cell-by-cell plots showing change in rise time, amplitude, decay time, and mean intercellular correlation following GABAergic block with bicuculline. Parameters were calculated between active cells. Marker color indicates cells from the same spheroid. Red dashed line indicates new mean, with 99% confidence interval shaded in on either side. Cells exhibited no significant change in mean rise time (t-test, p>0.01) but a modest increase in mean decay time (+7% ± 24%, p<0.01). Mean event amplitude decreased (−14% ± 18%, t-test, p<0.01) but intercellular correlation did not change (t-test, p>0.01). Cells shown in (A) are from spheroid #1. D) Amplitude increased and frequency decreased following GABAergic block with bicuculline in Day 14 spheroids. ΔF/F traces of cells in an 8K rat spheroid at baseline (left) and after GABAergic block with bicuculline (right). Red line represents 20 minute bicuculline incubation. Time series scale bars are 5 sec (horizontal) by 0.1 ΔF/F (vertical). E) Cells sometimes altered their calcium activity following GABAergic blockade with bicuculline at Day 14. Pie chart depicting proportions of cells that exhibited activity in both control and GABAergic blockade (violet, 57%) or never displayed dynamic calcium activity (blue, 25%). Cohorts of cells displayed activity solely under GABAergic blockade (green, 12%) or in the control condition (yellow, 6%). Results were taken from 163 cells across 8 spheres from 2 separate dissections. F) At Day 14, GABAergic blockade via bicuculline incubation yielded increases in correlation and decay time, and a decrease in rise time. Normalized cell-by-cell plots showing change in rise time, amplitude, decay time, and mean intercellular correlation following GABAergic block with bicuculline. Parameters were calculated between active cells. Marker color indicates cells from the same spheroid. Red dashed line indicates new mean correlation, with 99% confidence interval shaded in on either side. Cells exhibited a modest but variable decrease in mean rise time (−26% ± 82%, t-test, p>0.01) and an increase in mean decay time (+24% ± 58%, t-test, p<0.01). Mean event amplitude remained unchanged (t-test, p>0.01) but intercellular correlation increased dramatically (+48% ± 37%, t-test, p>0.01). Cells shown in (D) are from spheroid #2.

## Discussion

In the rapidly-evolving 3D *in vitro* culture landscape, increasing attention is being paid to the complex functional signatures of neuronal networks ^13^. We conducted a cross-sectional analysis of network development in postnatal rat cortical spheroids and found a progression of activity patterns. Some synchronized firing was observed in rat spheroids at Day 7, though this was highly variable. At Day 9, synchronized activity around 0.5 Hz became almost ubiquitous, with most and sometimes all the cells participating. Between Days 10 to 14, correlation decreased, giving way to sparse and complex patterns of activity. This activity did not statistically differ between male-derived and female-derived rat spheroids according to any of the metrics used.

Network activity did not appear to be influenced by sex, species, or type of calcium indicator. We explored excitatory (Fig 3G-I) and inhibitory (Fig 4) contributions to network dynamics, and demonstrated that these contributions change over time. Our scaffold-free, self-assembly approach leads to a spheroid that contains cortical extracellular matrix, stiffness, cell density, and cell types including neurons, astrocytes, microglia, and endothelial cells ^3^. The development of network oscillations in weeks rather than months is likely due to a combination of the presence of non-neuronal lineage cell types, as well as their differentiated state ^27^. This is supported by the similarly rapid time to activity generation in scaffold cultures of primary mouse neurons ^28^. The ready availability of rodent cortical cells – a neonatal rat brain yields five to nine hundred 8K spheroids, and a neonatal mouse brain yields about two to four hundred – lends itself to high throughput experiments in size-controlled 3D tissues. Rodent 3D culture models present an opportunity to strengthen *in vitro* to *in vivo* extrapolations based on a vast array of experiments, gaining the advantages of numerous *in vivo* behavioral assays in rats, as well as the powerful genetic tools available in mice. Because baseline activity was uniform across sex, differences in response to manipulations or therapeutics would reveal possible sexual dimorphisms, thus demonstrating the need for further study. This can potentially simplify the testing of therapeutics in both male and female models as the field strives to correct for the prior five-fold overrepresentation of male animals in neuroscience studies ^29^. This model also presents an opportunity to study diseases with disparate outcomes between the sexes.

The formation of neural network activity has been investigated in several types of *in vitro* systems, including 2D microelectrode array output of rat hippocampal neurons in glass microbead-poly dimethyl siloxane scaffolds ^30^; OGB-Ca imaging of adherent cell assemblies of human pluripotent stem cell-derived cerebral cortex neurons ^31^; 2D microelectrode array output of human cortical organoids ^2^; and Ca imaging of mouse GCaMP neurons in a silk scaffold ^28^ . Evidence suggests that 2D and 3D configurations produce different network structures ^30,32,33^, and that the 3D networks more closely approximate in vivo networks ^32,33^. The number of cortical cells in a given network affects the patterns of activity the network can produce. While it can be difficult to estimate the number of cells in iPSC-derived cultures due to varying differentiation rates, rodent cortical spheroids are seeded at a calculated density of cells which include reproducible populations of glia and differentiated neurons ^3^. In these experiments, networks of different sizes displayed different patterns and timelines of activity development. Activity patterns of 2K, 4K, and 8K spheroids were monitored over the first weeks of culture, with only 8K spheroids showing reproducible activity in the first weeks of culture. Interestingly, this latency to the development of network activity is comparable to what was found in primary mouse neurons grown in scaffolds ^28^, in spite of vastly different network sizes. It should be noted however, that the scale of activity under investigation in each paradigm was also different. In the case of a 2 million cell culture suspended in a silk or collagen scaffold, hexagonal ROIs of 500 μm diameter were created. The use herein of 8K rodent cortical spheroids allows spatiotemporally precise measurements (10-20μm ROIs) that allow for the investigation of more localized circuitry.

Functional networks require a balance of excitatory and inhibitory synapses; in the cortex, glutamatergic activity is spatiotemporally modulated by local GABAergic inhibition ^34^. Consistent with this, we showed that cortical spheroid activity was sensitive to both excitatory and inhibitory inputs. Elimination of glutamatergic inputs with NBQX/APV led to cessation of activity. Although GABAergic blockade by addition of bicuculline at Day 10 led to small decreases in amplitude, the same concentration of bicuculline at Day 14 evoked sharp increases in intracellular synchrony. Day 14 GABAergic blockade also yielded an increase in the duration of intracellular calcium signal, with mean rise time decreasing and mean decay time increasing. These results are consistent with the changing role of GABA_A_ receptors from excitatory to inhibitory signaling during development ^35,36^, although baseline increases in extracellular-GABA-mediated tonic inhibition may also play a role ^37^. Alteration of the excitatory/inhibitory balance in either direction led to detectable changes in network properties. We highlighted several fluorescence-based metrics by which to compare spheroids, offering future studies several metrics by which to measure differences between spheroids subjected to disparate conditions, or even spheroids derived from different genetic sources. Together, these results demonstrate that emergent network phenomena occur *in vitro*, and offer the potential for measuring these phenomena during network or tissue perturbations.

## Conclusions

In summary, we showed the development of network activity over the first weeks of 3D rodent cortical spheroid cultures. Calcium imaging at multiple time points showed development of complex network dynamics, transitioning from sporadic firing, to synchronized oscillations, to intermittent firing. These results create a framework for analyzing the functional outputs of neural cultures. We recorded from dozens of spheroids across species, sexes, and developmental timepoints. Pharmacological experiments underscore the sensitivity of these spheroids to disruptions in excitatory-inhibitory balance. Recording from groups of spheroids allows us to account for potential batch effects. Taken together, this work offers a robust baseline upon which to base future systems-level *in vitro* studies.

## Author Contributions

JLS and DHK conceptualized and designed the study. JLS and BT conducted experiments. JLS wrote custom software. JLS analyzed and visualized the data. DHK and BT provided necessary resources. JLS wrote the original draft. JLS, BT, and DHK edited and revised the paper.

## Conflicts of interest

There are no conflicts to declare.

## Acknowledgements

The authors thank Christien Hernandez and Lisa Osaki for help with data annotation, Harrison Katz for assistance with code and data analysis, and Rachel McLaughlin and Liane Livi for culturing assistance and helpful discussion. The authors also thank Jason Ritt for helpful discussion and critical reading of the manuscript.

This research was supported by a Brown University Presidential Fellowship (JLS), by BRP award NIEHS U01ES028184 (DHK), and by the U. S. Office of Naval Research under PANTHER award number N000142112044 (DHK) through Dr. Timothy Bentley.

## Notes

### Competing Interest Statement

The authors have declared no competing interest.

