## Supplementary Figures and Tables for "Cortical spheroids display oscillatory network dynamics"

**Electronic Supplementary Information:**

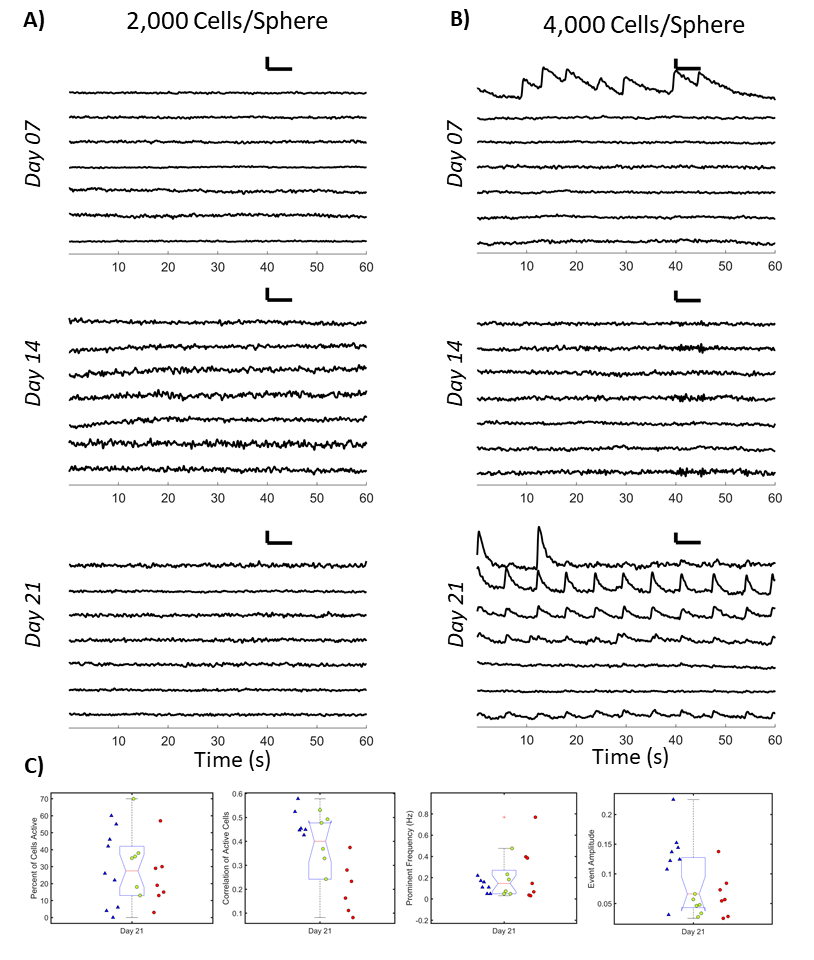

**Figure S1)** **Coordinated activity failed to develop in spheroids composed of 2,000 cells; emerged by Day 21 in 4K spheroids.** **A)** Activity was not detected in 2K spheroids. Representative ΔF/F traces from 2,000 cell spheroids at Day 7 (top), Day 14 (center) and Day 21 (bottom). Time series scale bars are 5 sec (horizontal) by 0.1 ΔF/F (vertical). **B)** Some sporadic activity was evident in 4K spheroids. Representative ΔF/F traces from 4K spheroids at Day 7 (top), Day 14 (center) and Day 21 (bottom). Time series scale bars are 5 sec (horizontal) by 0.1 ΔF/F (vertical). **C)** Percent of cells active, correlation of active cells, amplitude of events, and frequency of events were calculated for 4K spheroids imaged at Day 21 (n = 18). Marker symbol indicates the sex of rodent the spheroid was derived from (▲=♂, ●=♀). Marker color ranges from violet to maroon and indicates which data points came from the same dissections. t-tests without assuming equal variance were conducted and found no significant differences between means of male- and female-derived spheroids in any category with α=0.01. Red + symbols indicate statistically detected outliers.

**Table 2.S1)** Individual Data for Rat, 8K, Control/Spontaneous Condition.

| Date  YYMMDD | Dissection # | Species | Size | Sex | Days In Vitro | Sphere # | Avg Event Amplitude | Avg Event Rise Time (s) | Avg Event 50% Decay Time (s) | Percent of Cells Active | Correlation of Active Cells | Prevalent Frequency (Hz) |
| --- | --- | --- | --- | --- | --- | --- | --- | --- | --- | --- | --- | --- |
| 180417 | 1 | RT | 8K | F | 07 | 01 | 0.000 | 0.000 | 0.000 | 0 | 0.000 | 0.000 |
| 180417 | 1 | RT | 8K | F | 07 | 02 | 0.126 | 1.381 | 0.870 | 2 |  | 0.330 |
| 180417 | 1 | RT | 8K | F | 07 | 03 | 0.075 | 1.037 | 1.122 | 9 | 0.035 | 0.378 |
| 180417 | 1 | RT | 8K | F | 07 | 04 | 0.104 | 1.633 | 1.410 | 24 | 0.693 | 0.012 |
| 180417 | 1 | RT | 8K | F | 07 | 05 | 0.132 | 1.242 | 1.614 | 5 | -0.070 | 0.256 |
| 180417 | 1 | RT | 8K | F | 07 | 06 | 0.086 | 1.146 | 1.150 | 16 | 0.287 | 0.378 |
| 180417 | 1 | RT | 8K | F | 07 | 08 | 0.000 | 0.000 | 0.000 | 0 | 0.000 | 0.000 |
| 181118 | 6 | RT | 8K | F | 07 | 01 | 0.229 | 1.629 | 1.923 | 67 | 0.450 | 0.061 |
| 181118 | 6 | RT | 8K | F | 07 | 02 | 0.185 | 1.466 | 1.859 | 91 | 0.561 | 0.073 |
| 181118 | 6 | RT | 8K | F | 07 | 03 | 0.101 | 1.897 | 1.079 | 100 | 0.889 | 0.012 |
| 181118 | 6 | RT | 8K | F | 07 | 04 | 0.091 | 1.630 | 1.284 | 92 | 0.456 | 0.073 |
| 181118 | 6 | RT | 8K | F | 07 | 05 | 0.134 | 1.621 | 1.463 | 93 | 0.571 | 0.110 |
| 181118 | 6 | RT | 8K | F | 07 | 06 | 0.136 | 1.894 | 1.606 | 88 | 0.566 | 0.012 |
| 181118 | 6 | RT | 8K | F | 07 | 07 | 0.094 | 1.481 | 1.298 | 95 | 0.547 | 0.061 |
| 181118 | 6 | RT | 8K | F | 07 | 08 | 0.115 | 1.473 | 1.602 | 95 | 0.639 | 0.085 |
| 181204 | 7 | RT | 8K | F | 07 | 01 | 0.048 | 1.019 | 0.925 | 53 | 0.110 | 0.500 |
| 181204 | 7 | RT | 8K | F | 07 | 02 | 0.039 | 1.110 | 0.872 | 67 | 0.280 | 0.427 |
| 181204 | 7 | RT | 8K | F | 07 | 03 | 0.060 | 1.120 | 0.967 | 91 | 0.231 | 0.378 |
| 181204 | 7 | RT | 8K | F | 07 | 04 | 0.045 | 0.917 | 0.929 | 75 | 0.369 | 0.439 |
| 181204 | 7 | RT | 8K | F | 07 | 05 | 0.045 | 0.970 | 0.823 | 59 | 0.396 | 0.452 |
| 181204 | 7 | RT | 8K | F | 07 | 06 | 0.051 | 1.209 | 1.088 | 53 | 0.084 | 0.061 |
| 181204 | 7 | RT | 8K | F | 07 | 07 | 0.038 | 1.158 | 1.016 | 59 | 0.155 | 0.366 |
| 181204 | 7 | RT | 8K | F | 07 | 08 | 0.046 | 1.195 | 1.069 | 44 | 0.106 | 0.171 |
| 180417 | 1 | RT | 8K | M | 07 | 01 | 0.092 | 1.083 | 1.918 | 4 | 0.156 | 0.159 |
| 180417 | 1 | RT | 8K | M | 07 | 02 | 0.118 | 0.964 | 1.226 | 20 | 0.384 | 0.232 |
| 180417 | 1 | RT | 8K | M | 07 | 03 | 0.205 | 1.720 | 1.197 | 4 | 0.013 | 0.110 |
| 180417 | 1 | RT | 8K | M | 07 | 04 | 0.176 | 1.123 | 1.438 | 14 | -0.018 | 0.146 |
| 180417 | 1 | RT | 8K | M | 07 | 07 | 0.114 | 1.530 | 1.325 | 11 | -0.019 | 0.012 |
| 180417 | 1 | RT | 8K | M | 07 | 08 | 0.172 | 1.234 | 2.116 | 13 | -0.030 | 0.159 |
| 181118 | 6 | RT | 8K | M | 07 | 01 | 0.061 | 1.134 | 0.876 | 89 | 0.479 | 0.293 |
| 181118 | 6 | RT | 8K | M | 07 | 02 | 0.082 | 1.150 | 1.011 | 65 | 0.418 | 0.256 |
| 181118 | 6 | RT | 8K | M | 07 | 03 | 0.072 | 1.168 | 1.078 | 88 | 0.445 | 0.244 |
| 181118 | 6 | RT | 8K | M | 07 | 04 | 0.062 | 1.321 | 1.044 | 91 | 0.552 | 0.183 |
| 181118 | 6 | RT | 8K | M | 07 | 05 | 0.077 | 1.323 | 1.058 | 82 | 0.357 | 0.195 |
| 181118 | 6 | RT | 8K | M | 07 | 06 | 0.097 | 1.239 | 1.044 | 92 | 0.612 | 0.208 |
| 190212 | 8 | RT | 8K | M | 07 | 03 | 0.043 | 1.199 | 1.040 | 42 | 0.258 | 0.378 |
| 190212 | 8 | RT | 8K | M | 07 | 04 | 0.048 | 0.965 | 0.757 | 40 | 0.091 | 0.391 |
| 190212 | 8 | RT | 8K | M | 07 | 05 | 0.053 | 1.055 | 0.861 | 68 | 0.615 | 0.305 |
| 190212 | 8 | RT | 8K | M | 07 | 06 | 0.043 | 1.150 | 0.903 | 47 | 0.314 | 0.317 |
| 190212 | 8 | RT | 8K | M | 07 | 07 | 0.055 | 1.157 | 0.937 | 73 | 0.295 | 0.305 |
| 180420 | 1 | RT | 8K | F | 09 | 01 | 0.031 | 0.804 | 0.714 | 3 |  | 0.146 |
| 180420 | 1 | RT | 8K | F | 09 | 02 | 0.156 | 1.244 | 1.057 | 61 | 0.660 | 0.220 |
| 180420 | 1 | RT | 8K | F | 09 | 03 | 0.075 | 0.903 | 0.824 | 9 | 0.150 | 0.525 |
| 180420 | 1 | RT | 8K | F | 09 | 04 | 0.198 | 1.342 | 1.229 | 88 | 0.570 | 0.134 |
| 180420 | 1 | RT | 8K | F | 09 | 05 | 0.183 | 1.183 | 1.092 | 64 | 0.665 | 0.195 |
| 180420 | 1 | RT | 8K | F | 09 | 06 | 0.148 | 1.391 | 1.320 | 80 | 0.641 | 0.110 |
| 180420 | 1 | RT | 8K | F | 09 | 07 | 0.109 | 1.262 | 1.341 | 72 | 0.635 | 0.195 |
| 180420 | 1 | RT | 8K | F | 09 | 08 | 0.108 | 1.310 | 1.349 | 71 | 0.650 | 0.195 |
| 180913 | 3 | RT | 8K | F | 09 | 01 | 0.079 | 0.870 | 0.710 | 86 | 0.533 | 0.513 |
| 180913 | 3 | RT | 8K | F | 09 | 01 | 0.079 | 0.870 | 0.710 | 86 | 0.533 | 0.513 |
| 180913 | 3 | RT | 8K | F | 09 | 02 | 0.141 | 0.955 | 0.818 | 94 | 0.720 | 0.293 |
| 180913 | 3 | RT | 8K | F | 09 | 03 | 0.093 | 1.001 | 0.838 | 93 | 0.554 | 0.366 |
| 180913 | 3 | RT | 8K | F | 09 | 04 | 0.121 | 0.915 | 0.707 | 94 | 0.773 | 0.391 |
| 180913 | 3 | RT | 8K | F | 09 | 05 | 0.156 | 0.984 | 0.771 | 100 | 0.787 | 0.305 |
| 180913 | 3 | RT | 8K | F | 09 | 06 | 0.062 | 0.982 | 0.670 | 94 | 0.687 | 0.366 |
| 180913 | 3 | RT | 8K | F | 09 | 07 | 0.075 | 0.994 | 0.694 | 100 | 0.750 | 0.354 |
| 180913 | 3 | RT | 8K | F | 09 | 08 | 0.067 | 0.904 | 0.723 | 100 | 0.790 | 0.366 |
| 180927 | 4 | RT | 8K | F | 09 | 02 | 0.117 | 0.874 | 0.695 | 56 | 0.490 | 0.427 |
| 180927 | 4 | RT | 8K | F | 09 | 04 | 0.234 | 1.453 | 1.006 | 97 | 0.703 | 0.171 |
| 180927 | 4 | RT | 8K | F | 09 | 05 | 0.072 | 0.917 | 0.809 | 75 | 0.464 | 0.476 |
| 180927 | 4 | RT | 8K | F | 09 | 11 | 0.130 | 0.874 | 0.798 | 87 | 0.617 | 0.464 |
| 180927 | 4 | RT | 8K | F | 09 | 13 | 0.069 | 0.889 | 0.717 | 76 | 0.489 | 0.452 |
| 180927 | 4 | RT | 8K | F | 09 | 15 | 0.075 | 0.964 | 0.677 | 83 | 0.568 | 0.452 |
| 181120 | 6 | RT | 8K | F | 09 | 01 | 0.046 | 0.905 | 0.638 | 92 | 0.537 | 0.549 |
| 181120 | 6 | RT | 8K | F | 09 | 02 | 0.048 | 0.799 | 0.573 | 73 | 0.585 | 0.635 |
| 181120 | 6 | RT | 8K | F | 09 | 03 | 0.051 | 0.861 | 0.579 | 96 | 0.583 | 0.562 |
| 181120 | 6 | RT | 8K | F | 09 | 04 | 0.041 | 0.879 | 0.633 | 89 | 0.461 | 0.623 |
| 181120 | 6 | RT | 8K | F | 09 | 06 | 0.048 | 0.928 | 0.642 | 80 | 0.531 | 0.537 |
| 190427 | 9 | RT | 8K | F | 09 | 01 | 0.034 | 1.061 | 0.497 | 5 | 0.286 | 0.940 |
| 190427 | 9 | RT | 8K | F | 09 | 02 | 0.050 | 1.026 | 0.580 | 33 | 0.270 | 0.659 |
| 190427 | 9 | RT | 8K | F | 09 | 03 | 0.044 | 1.161 | 0.602 | 24 | 0.323 | 0.610 |
| 190427 | 9 | RT | 8K | F | 09 | 04 | 0.046 | 1.130 | 0.672 | 45 | 0.351 | 0.574 |
| 190427 | 9 | RT | 8K | F | 09 | 05 | 0.045 | 0.982 | 0.606 | 38 | 0.546 | 0.549 |
| 190427 | 9 | RT | 8K | F | 09 | 06 | 0.044 | 1.006 | 0.576 | 40 | 0.429 | 0.549 |
| 190427 | 9 | RT | 8K | F | 09 | 08 | 0.047 | 1.035 | 0.579 | 49 | 0.465 | 0.537 |
| 190815 | 11 | RT | 8K | F | 09 | 01 | 0.077 | 0.914 | 0.642 | 47 | 0.497 | 0.415 |
| 190815 | 11 | RT | 8K | F | 09 | 04 | 0.086 | 1.036 | 0.747 | 52 | 0.434 | 0.073 |
| 190815 | 11 | RT | 8K | F | 09 | 05 | 0.064 | 0.774 | 0.527 | 56 | 0.736 | 0.525 |
| 190815 | 11 | RT | 8K | F | 09 | 06 | 0.068 | 0.946 | 0.614 | 58 | 0.553 | 0.500 |
| 190815 | 11 | RT | 8K | F | 09 | 02 | 0.132 | 0.766 | 0.704 | 30 | 0.679 | 0.415 |
| 190815 | 11 | RT | 8K | F | 09 | 03 | 0.088 | 0.807 | 0.580 | 81 | 0.719 | 0.415 |
| 180420 | 1 | RT | 8K | M | 09 | 02 | 0.186 | 0.970 | 1.012 | 38 | 0.332 | 0.330 |
| 180420 | 1 | RT | 8K | M | 09 | 03 | 0.126 | 0.863 | 0.816 | 44 | 0.493 | 0.378 |
| 180420 | 1 | RT | 8K | M | 09 | 04 | 0.118 | 1.216 | 1.018 | 33 | 0.351 | 0.269 |
| 180420 | 1 | RT | 8K | M | 09 | 05 | 0.103 | 1.219 | 1.124 | 32 | -0.019 | 0.012 |
| 180420 | 1 | RT | 8K | M | 09 | 06 | 0.144 | 1.213 | 1.491 | 52 | 0.501 | 0.122 |
| 180420 | 1 | RT | 8K | M | 09 | 07 | 0.106 | 1.341 | 0.993 | 48 | 0.558 | 0.281 |
| 180420 | 1 | RT | 8K | M | 09 | 08 | 0.117 | 1.151 | 0.987 | 62 | 0.513 | 0.281 |
| 180913 | 3 | RT | 8K | M | 09 | 01 | 0.077 | 0.858 | 0.710 | 52 | 0.474 | 0.574 |
| 180913 | 3 | RT | 8K | M | 09 | 02 | 0.082 | 0.885 | 0.639 | 97 | 0.729 | 0.500 |
| 180913 | 3 | RT | 8K | M | 09 | 03 | 0.044 | 0.691 | 0.686 | 36 | 0.247 | 0.574 |
| 180913 | 3 | RT | 8K | M | 09 | 04 | 0.051 | 0.969 | 0.702 | 34 | 0.225 | 0.647 |
| 180913 | 3 | RT | 8K | M | 09 | 05 | 0.061 | 1.017 | 0.800 | 98 | 0.715 | 0.305 |
| 180913 | 3 | RT | 8K | M | 09 | 06 | 0.060 | 0.951 | 0.706 | 84 | 0.606 | 0.403 |
| 180913 | 3 | RT | 8K | M | 09 | 07 | 0.067 | 0.969 | 0.829 | 87 | 0.734 | 0.293 |
| 180913 | 3 | RT | 8K | M | 09 | 08 | 0.044 | 1.046 | 0.784 | 83 | 0.525 | 0.378 |
| 190927 | 12 | RT | 8K | M | 09 | 07 | 0.246 | 1.334 | 1.157 | 75 | 0.685 | 0.244 |
| 190927 | 12 | RT | 8K | M | 09 | 08 | 0.103 | 1.184 | 0.834 | 69 | 0.593 | 0.256 |
| 190927 | 12 | RT | 8K | M | 09 | 09 | 0.117 | 1.038 | 0.860 | 80 | 0.750 | 0.256 |
| 190927 | 12 | RT | 8K | M | 09 | 10 | 0.112 | 1.160 | 0.879 | 76 | 0.657 | 0.220 |
| 190215 | 8 | RT | 8K | F | 10 | 01 | 0.046 | 1.108 | 0.915 | 42 | 0.328 | 0.391 |
| 190215 | 8 | RT | 8K | F | 10 | 02 | 0.052 | 1.111 | 0.890 | 23 | 0.213 | 0.391 |
| 190215 | 8 | RT | 8K | F | 10 | 03 | 0.047 | 1.220 | 0.923 | 51 | 0.318 | 0.317 |
| 190215 | 8 | RT | 8K | F | 10 | 04 | 0.059 | 1.091 | 1.006 | 48 | 0.212 | 0.305 |
| 190215 | 8 | RT | 8K | F | 10 | 05 | 0.048 | 0.966 | 0.816 | 57 | 0.499 | 0.330 |
| 190215 | 8 | RT | 8K | F | 10 | 07 | 0.052 | 1.025 | 0.974 | 62 | 0.330 | 0.342 |
| 180817 | 2 | RT | 8K | M | 10 | 01 | 0.092 | 0.615 | 0.672 | 9 | 0.537 | 0.745 |
| 180817 | 2 | RT | 8K | M | 10 | 09 | 0.068 | 0.884 | 0.875 | 21 | 0.128 | 0.586 |
| 180817 | 2 | RT | 8K | M | 10 | 10 | 0.093 | 0.830 | 0.808 | 27 | 0.277 | 0.586 |
| 180817 | 2 | RT | 8K | M | 10 | 11 | 0.076 | 0.735 | 0.556 | 15 | 0.396 | 0.610 |
| 180817 | 2 | RT | 8K | M | 10 | 12 | 0.075 | 0.782 | 0.901 | 36 | 0.213 | 0.525 |
| 180817 | 2 | RT | 8K | M | 10 | 13 | 0.082 | 0.772 | 0.791 | 32 | 0.425 | 0.623 |
| 180817 | 2 | RT | 8K | M | 10 | 14 | 0.061 | 0.802 | 0.648 | 25 | 0.431 | 0.806 |
| 180817 | 2 | RT | 8K | M | 10 | 15 | 0.065 | 0.815 | 1.126 | 33 | 0.243 | 0.549 |
| 180817 | 2 | RT | 8K | M | 10 | 16 | 0.082 | 0.675 | 0.728 | 40 | 0.686 | 0.500 |
| 190215 | 8 | RT | 8K | M | 10 | 01 | 0.039 | 1.056 | 0.872 | 64 | 0.293 | 0.488 |
| 190215 | 8 | RT | 8K | M | 10 | 05 | 0.048 | 1.116 | 0.862 | 70 | 0.527 | 0.354 |
| 190215 | 8 | RT | 8K | M | 10 | 07 | 0.040 | 0.998 | 0.908 | 77 | 0.358 | 0.476 |
| 180424 | 1 | RT | 8K | F | 14 | 01 | 0.130 | 1.280 | 0.786 | 46 | 0.508 | 0.159 |
| 180424 | 1 | RT | 8K | F | 14 | 02 | 0.132 | 1.374 | 0.704 | 30 | 0.279 | 0.122 |
| 180424 | 1 | RT | 8K | F | 14 | 03 | 0.122 | 0.957 | 0.748 | 21 | 0.082 | 0.500 |
| 180424 | 1 | RT | 8K | F | 14 | 04 | 0.141 | 1.199 | 0.678 | 28 | 0.314 | 0.171 |
| 180424 | 1 | RT | 8K | F | 14 | 05 | 0.118 | 1.134 | 0.573 | 17 | 0.246 | 0.586 |
| 180424 | 1 | RT | 8K | F | 14 | 06 | 0.125 | 1.308 | 1.121 | 27 | 0.418 | 0.256 |
| 180424 | 1 | RT | 8K | F | 14 | 07 | 0.081 | 0.888 | 0.571 | 36 | 0.541 | 0.598 |
| 180424 | 1 | RT | 8K | F | 14 | 08 | 0.106 | 1.002 | 0.575 | 28 | 0.140 | 0.452 |
| 181113 | 5 | RT | 8K | F | 14 | 01 | 0.101 | 0.803 | 0.714 | 5 | -0.019 | 0.659 |
| 181113 | 5 | RT | 8K | F | 14 | 03 | 0.077 | 0.521 | 0.438 | 5 | 0.023 | 0.610 |
| 181113 | 5 | RT | 8K | F | 14 | 04 | 0.072 | 0.762 | 1.078 | 9 | 0.000 | 0.061 |
| 181113 | 5 | RT | 8K | F | 14 | 05 | 0.097 | 0.643 | 0.752 | 5 | 0.111 | 0.549 |
| 181113 | 5 | RT | 8K | F | 14 | 06 | 0.111 | 1.639 | 2.328 | 76 | 0.941 | 0.256 |
| 181113 | 5 | RT | 8K | F | 14 | 07 | 0.043 | 0.679 | 0.823 | 5 | 0.059 | 0.378 |
| 181113 | 5 | RT | 8K | F | 14 | 08 | 0.105 | 0.774 | 0.795 | 12 | 0.029 | 0.464 |
| 181211 | 7 | RT | 8K | F | 14 | 01 | 0.179 | 0.852 | 1.147 | 88 | 0.792 | 0.244 |
| 181211 | 7 | RT | 8K | F | 14 | 02 | 0.181 | 0.898 | 1.101 | 87 | 0.733 | 0.232 |
| 181211 | 7 | RT | 8K | F | 14 | 03 | 0.099 | 0.893 | 1.120 | 85 | 0.713 | 0.281 |
| 181211 | 7 | RT | 8K | F | 14 | 04 | 0.051 | 0.918 | 0.704 | 94 | 0.470 | 0.549 |
| 181211 | 7 | RT | 8K | F | 14 | 05 | 0.069 | 0.897 | 0.677 | 78 | 0.515 | 0.464 |
| 181211 | 7 | RT | 8K | F | 14 | 06 | 0.069 | 1.004 | 1.010 | 94 | 0.574 | 0.256 |
| 181211 | 7 | RT | 8K | F | 14 | 07 | 0.069 | 1.032 | 0.808 | 96 | 0.486 | 0.293 |
| 181211 | 7 | RT | 8K | F | 14 | 08 | 0.049 | 1.091 | 0.803 | 100 | 0.336 | 0.500 |
| 190626 | 10 | RT | 8K | F | 14 | 01 | 0.219 | 0.959 | 1.164 | 6 | -0.340 | 0.403 |
| 190626 | 10 | RT | 8K | F | 14 | 04 | 0.085 | 1.250 | 0.567 | 7 | -0.014 | 0.427 |
| 190626 | 10 | RT | 8K | F | 14 | 06 | 0.099 | 1.175 | 0.656 | 49 | 0.459 | 0.476 |
| 190626 | 10 | RT | 8K | F | 14 | 07 | 0.117 | 1.001 | 0.781 | 32 | 0.463 | 0.562 |
| 180821 | 2 | RT | 8K | M | 14 | 01 | 0.090 | 0.614 | 0.523 | 2 | 0.000 | 0.049 |
| 180821 | 2 | RT | 8K | M | 14 | 02 | 0.103 | 1.148 | 1.097 | 6 | -0.091 | 0.049 |
| 180821 | 2 | RT | 8K | M | 14 | 03 | 0.224 | 2.858 | 1.406 | 13 | -0.087 | 0.049 |
| 180821 | 2 | RT | 8K | M | 14 | 04 | 0.035 | 0.859 | 0.644 | 6 | -0.004 | 0.061 |
| 180821 | 2 | RT | 8K | M | 14 | 05 | 0.041 | 0.879 | 0.766 | 19 | 0.015 | 0.110 |
| 180821 | 2 | RT | 8K | M | 14 | 06 | 0.077 | 2.169 | 1.585 | 13 | 0.022 | 0.061 |
| 180821 | 2 | RT | 8K | M | 14 | 07 | 0.043 | 0.858 | 0.782 | 9 | 0.007 | 0.110 |
| 180821 | 2 | RT | 8K | M | 14 | 08 | 0.059 | 0.965 | 0.673 | 10 | 0.004 | 0.159 |
| 180821 | 2 | RT | 8K | M | 14 | 10 | 0.094 | 1.262 | 0.992 | 15 | 0.109 | 0.134 |

**Table 2.S2)** Individual Data for Mouse, 8K, Control/GCaMP condition and OBG, DMSO conditions

| Date  YYMMDD | Species | Size | Sex | Days In Vitro | Sphere # | Condition | Avg Event Amplitude | Avg Event Rise Time (s) | Avg Event 50% Decay Time (s) | Percent of Cells Active | Correlation of Active Cells | Prominent Frequency |
| --- | --- | --- | --- | --- | --- | --- | --- | --- | --- | --- | --- | --- |
| 191126 | MS | 8K | M | 08 | 01 | GCAMP | 0.748 | 1.577 | 0.890 | 100 | 0.710 | 0.146 |
| 191126 | MS | 8K | M | 08 | 02 | GCAMP | 1.029 | 5.361 | 1.088 | 95 | 0.867 | 0.073 |
| 191126 | MS | 8K | M | 08 | 04 | GCAMP | 0.503 | 1.121 | 0.777 | 93 | 0.498 | 0.232 |
| 191126 | MS | 8K | M | 08 | 05 | GCAMP | 0.870 | 1.577 | 0.936 | 100 | 0.488 | 0.122 |
| 191126 | MS | 8K | M | 08 | 06 | GCAMP | 0.744 | 1.610 | 0.835 | 94 | 0.645 | 0.012 |
| 191126 | MS | 8K | M | 08 | 01 | DMSOF127 | 0.864 | 1.230 | 0.671 | 92 | 0.347 | 0.110 |
| 191126 | MS | 8K | M | 08 | 02 | DMSOF127 | 0.540 | 1.515 | 0.636 | 100 | 0.268 | 0.159 |
| 191126 | MS | 8K | M | 08 | 03 | DMSOF127 | 0.360 | 1.242 | 0.528 | 95 | 0.187 | 0.085 |
| 191126 | MS | 8K | M | 08 | 04 | DMSOF127 | 0.131 | 1.409 | 0.739 | 93 | 0.516 | 0.073 |
| 191126 | MS | 8K | M | 08 | 05 | DMSOF127 | 0.753 | 0.906 | 0.593 | 100 | 0.919 | 0.256 |
| 191126 | MS | 8K | M | 08 | 01 | OGB | 0.117 | 0.978 | 0.586 | 96 | 0.513 | 0.415 |
| 191126 | MS | 8K | M | 08 | 02 | OGB | 0.068 | 1.087 | 0.712 | 91 | 0.470 | 0.342 |
| 191126 | MS | 8K | M | 08 | 03 | OGB | 0.066 | 1.068 | 0.667 | 76 | 0.367 | 0.403 |
| 191201 | MS | 8K | M | 13 | 03 | GCAMP | 0.605 | 1.438 | 0.770 | 100 | 0.587 | 0.171 |
| 191201 | MX | 8K | M | 13 | 05 | GCAMP | 0.534 | 0.886 | 0.575 | 94 | 0.738 | 0.073 |
| 191201 | MS | 8K | M | 13 | 06 | GCAMP | 0.861 | 1.469 | 0.894 | 100 | 0.857 | 0.122 |
| 191201 | MS | 8K | M | 13 | 07 | GCAMP | 1.099 | 1.026 | 0.903 | 100 | 0.956 | 0.183 |
| 191201 | MS | 8K | M | 13 | 08 | GCAMP | 0.772 | 1.809 | 1.384 | 92 | 0.856 | 0.110 |
| 191201 | MS | 8K | M | 13 | 14 | GCAMP | 1.447 | 2.350 | 0.840 | 100 | 0.883 | 0.098 |
| 191201 | MS | 8K | M | 13 | 15 | GCAMP | 0.677 | 1.770 | 0.515 | 100 | 0.559 | 0.183 |
| 191201 | MS | 8K | M | 13 | 16 | GCAMP | 0.601 | 1.490 | 0.672 | 100 | 0.524 | 0.208 |
| 191201 | MS | 8K | M | 13 | 17 | GCAMP | 0.580 | 1.269 | 0.627 | 83 | 0.544 | 0.061 |

**Table 2.S3)** Individual Data for Rat, 4K, Day 21, Control/Spontaneous Condition.

| Date  YYMMDD | Diss-ection # | Species | Size | Sex | Days In Vitro | Sphere # | Recording Duration (s) | Avg Event Amplitude | Avg Event Rise Time (s) | Avg Event 50% Decay Time (s) | Percent of Cells Active | Correlation of Active Cells | Prominent Frequency |
| --- | --- | --- | --- | --- | --- | --- | --- | --- | --- | --- | --- | --- | --- |
| 180501 | 1 | RT | 4K | M | 21 | 01 | 60 | 0.108 | 1.245 | 1.107 | 26 | 0.524 | 0.220 |
| 180501 | 1 | RT | 4K | M | 21 | 02 | 60 | 0.031 | 1.071 | 0.928 | 4 |  | 7.556 |
| 180501 | 1 | RT | 4K | M | 21 | 03 | 60 | 0.122 | 1.407 | 1.186 | 42 | 0.578 | 0.171 |
| 180501 | 1 | RT | 4K | M | 21 | 04 | 60 | 0.137 | 1.744 | 1.468 | 46 | 0.447 | 0.110 |
| 180501 | 1 | RT | 4K | M | 21 | 04 | 60 | 0.225 | 1.433 | 1.284 | 60 | 0.454 | 0.159 |
| 180501 | 1 | RT | 4K | M | 21 | 05 | 60 |  |  |  | 0 |  |  |
| 180501 | 1 | RT | 4K | M | 21 | 06 | 60 | 0.152 | 1.136 | 0.703 | 22 | 0.426 | 0.049 |
| 180501 | 1 | RT | 4K | M | 21 | 07 | 60 | 0.144 | 1.423 | 0.946 | 55 | 0.449 | 0.110 |
| 180501 | 1 | RT | 4K | M | 21 | 08 | 60 | 0.124 | 1.203 | 0.760 | 6 |  | 0.049 |
| 191120 | 13 | RT | 4K | F | 21 | 05 | 60 | 0.057 | 1.372 | 0.750 | 35 | 0.532 | 0.049 |
| 191120 | 13 | RT | 4K | F | 21 | 06 | 60 | 0.066 | 1.253 | 0.773 | 70 | 0.477 | 0.073 |
| 191120 | 13 | RT | 4K | F | 21 | 07 | 60 | 0.046 | 1.172 | 0.706 | 36 | 0.369 | 0.232 |
| 191120 | 13 | RT | 4K | F | 21 | 08 | 60 | 0.028 | 1.258 | 0.796 | 18 | 0.329 | 0.183 |
| 191120 | 13 | RT | 4K | F | 21 | 09 | 60 | 0.048 | 1.850 | 1.053 | 38 | 0.242 | 0.049 |
| 191120 | 13 | RT | 4K | F | 21 | 10 | 60 | 0.034 | 1.067 | 0.864 | 13 | 0.493 | 0.476 |
| 200106 | 14 | RT | 4K | F | 21 | 01 | 120 | 0.138 | 0.849 | 0.405 | 3 |  | 0.397 |
| 200106 | 14 | RT | 4K | F | 21 | 02 | 120 | 0.073 | 1.121 | 0.552 | 29 | 0.163 | 0.385 |
| 200106 | 14 | RT | 4K | F | 21 | 03 | 120 | 0.054 | 1.273 | 0.708 | 19 | 0.280 | 0.037 |
| 200106 | 14 | RT | 4K | F | 21 | 03 | 120 | 0.025 | 0.984 | 0.654 | 13 | 0.111 | 0.031 |
| 200106 | 14 | RT | 4K | F | 21 | 04 | 120 | 0.057 | 1.557 | 0.771 | 57 | 0.375 | 0.146 |
| 200106 | 14 | RT | 4K | F | 21 | 04 | 120 | 0.084 | 1.476 | 0.825 | 30 | 0.233 | 0.067 |
| 200106 | 14 | RT | 4K | F | 21 | 06 | 120 | 0.028 | 1.007 | 0.426 | 15 | 0.082 | 0.769 |
